# It’s not just poverty: unregulated global market and bad governance explain unceasing deforestation in Western Madagascar

**DOI:** 10.1101/2020.07.30.229104

**Authors:** Ghislain Vieilledent, Marie Nourtier, Clovis Grinand, Miguel Pedrono, Alison Clausen, Tsiky Rabetrano, Jean-Roger Rakotoarijaona, Bruno Rakotoarivelo, Fety A. Rakotomalala, Linjanantenaina Rakotomalala, Andriamandimbisoa Razafimpahanana, José M. Ralison, Frédéric Achard

## Abstract

Madagascar is recognized both for its unparalleled biodiversity and the high level of threat suffered by this biodiversity, associated in particular with anthropogenic deforestation. Despite sustained efforts to fight poverty and curb deforestation, forest cover in Madagascar is rapidly decreasing. To try to explain why it is so difficult to stop deforestation in Madagascar, we analysed the recent deforestation process in Western Madagascar through satellite image analysis and field surveys. We show that deforestation has increased from less than 0.9%/yr on 2000–2010 to more than 2%/yr on 2010–2017. We identified two major causes of deforestation, which were not associated with subsistence agriculture: slash-and-burn agriculture for the cultivation of cash crops (maize and peanut), and uncontrolled fires to create open pasture. Maize production is mainly at the destination of the domestic market and is used in particular for livestock feeding. Peanut production has boomed since 2013 and more than half of it is now exported towards asiatic countries. The money earned by farmers is principally invested into zebu herd acquisition. Trade of agricultural commodities benefits several intermediaries, some of whom have political responsibilities thus creating conflicts of interest. On the other hand, agents from institutions in charge of the management of the protected areas have no means to enforce laws against deforestation. In the absence of an efficient strategy to stop deforestation, we predicted that 38-93% of the forest present in 2000 will have disappeared in 2050. Forest loss, apart from biodiversity loss and climate-change global issues, will be at the expense of local population. In order to stop deforestation, international aid should be used to improve local governance to enforce environmental laws and pressure should be put on trading companies to buy certified agricultural commodities that are not derived from deforestation.

## 1 Introduction

Tropical forests provide important ecosystem services at the global scale, such as biodiversity conservation and climate regulation (Costanza *et al*., 1997), and at the local scale for peoples’ livelihoods (Anderson *et al*., 2006; Jeannoda *et al*., 2007; Gardner & Davies, 2014). On the island of Madagascar, three types of tropical forests, covering about 15% of the country area, can be found: the moist forest in the East, the xerophytic forest in the South, and the dry forest in the West (Humbert, 1955). Among all the ecosystem services they provide, Malagasy forests are particularly important for the unique biodiversity they shelter, both in terms of species diversity and endemism in many taxonomic groups (Goodman & Benstead, 2005; Brooks *et al*., 2006).

However, a large part of the original tropical forest is thought to have disappeared since the arrival of the humans on the island around 2000 years ago (Green & Sussman, 1990; Harper *et al*., 2007; Vieilledent *et al*., 2018b), some studies suggesting that 90% of the island was originally covered by forest (Burns *et al*., 2016). From 1953 to 2014, a loss of about 46% of the forest cover has been estimated (Vieilledent *et al*., 2018b), leading to an estimated extinction of 9% of the species between 1950 and 2000 in Madagascar (Allnutt *et al*., 2008). The common narrative attributes deforestation to the extreme poverty of the country and subsistence farming through slash-and-burn agriculture (Jarosz, 1993; Scales, 2011; Gardner *et al*., 2013). In 2010, about 81% of the population lived below the international poverty line of $1.25 per day. In 2015, more than 70% of the 24 million population of Madagascar (United Nations, 2017) rely heavily on forests for their livelihood (Anderson *et al*., 2006). Moreover, the Malagasy population is rapidly increasing, with population growing at a rate close to 3%/yr, resulting in a doubling each 25 years.

To curb deforestation, several conservation and rural development programs have been implemented in Madagascar since the establishment of the first protected areas in 1927. The Madagascar Protected Area System (SAPM) has seen both an increase in the number of protected areas and an increase in the place taken by rural development programs to accompany conservation actions (Virah-Sawmy *et al*., 2014). Rural development programs aimed at alleviating poverty, increasing agriculture productivity, and supporting education, health and local governance in the periphery of the protected areas (Freudenberger, 2010). Since 2003, Madagascar’s protected area coverage expanded from 1.7 to 6.4 Mha (11% of the country area, see Kremen *et al*. (2008) and Corson (2014)). In parallel with the development of the SAPM, the REDD+ (Reducing Emissions from Deforestation and Forest Degradation) mechanism have emerged in Madagascar to avoid deforestation combining conservation and development actions (Ebeling & Yasue, 2008). Five REDD+ pilot projects have been initiated in Madagascar since 2004 over a total of 1.76 Mha of forest (Demaze, 2014).

All of these actions were funded by the donor community, with some limited revenues from tourism entry fees to national parks. Donor efforts were led by the World Bank and USAID (United States Agency for International Development) (Freudenberger, 2010). The Malagasy Government does not contribute financially to the management of the protected area network. The total funding of the National Environmental Action Program (NEAP) from 1990 to 2010 was estimated at approximately $450 million for environment activities, with another 50% for related development programs (e.g. agriculture and health interventions) (Freudenberger, 2010). Other bilateral donors (mainly Europe, France and Germany) have also funded specific socio-environmental projects ($102 million in the period 2005–2011) (Critical Ecosystem Partnership Fund, 2014). The main international NGOs (World Wide Fund for Nature, Conservation International, Wildlife Conservation Society) and international research centers (such as CIRAD and IRD French research centers) also contributed significantly to the financial effort by securing alternative resources to support conservation and development programs (Critical Ecosystem Partnership Fund, 2014). Regarding REDD+ activities, the private sector also provided substantial funding. Air France supported the first phase of the PHCF REDD+ project (2008–2012) with $5 million and the Makira REDD+ project managed by the Wildlife Conservation Society made close to $3 million in sales between 2013 and 2018. Annually, this represents a rough estimate of about $60 million per year from the donor community to fund environmental and related development activities in Madagascar (80% of the annual national budget for the environment sector) (Critical Ecosystem Partnership Fund, 2014).

Despite all the efforts and money invested in conservation and development programs over the past 30 years (1985–2015), deforestation has not stopped on the island. Out of the 10.8 Mha of tropical forest in 1990, around 1.5 Mha (13% of the forest) was deforested during the period 1990–2010 (Harper *et al*., 2007; Vieilledent *et al*., 2018b), which corresponds to an average annual deforestation rate of 75,000 ha/yr (0.72%/yr). More worryingly, Madagascar has seen an acceleration of the deforestation recently (Vieilledent *et al*., 2018b). Madagascar was the third country in the world with the fastest acceleration of tree cover loss (+8.3% of deforested area per year) in the period 2001–2014 (see Hansen *et al*. (2013) and http://ow.ly/RBbgw). In the period 2010–2015, deforestation reached an average rate of 110,000 ha/yr (1.2%/yr). Forest cover in 2015 has been estimated at 8.8 Mha (Vieilledent *et al*., 2018b,a). The conclusion is unfortunately rather clear: conservation and development programs since the mid 1980s have failed to save Madagascar’s forests. It is therefore legitimate to wonder why such initiatives have not been successful and what could be alternative solutions to stop deforestation. Also, the recent deforestation might be attributable to new factors, independent of local human activities, such as global climate change, cyclone and natural fires, which would imply the need for conservation measures different to the ones used in the past.

To try to answer these questions, we undertook an in-depth analysis of the deforestation process in three study areas in Western Madagascar. The study areas were located around three protected areas: the Menabe-Antimena New Protected Area (acronym MANAP in French), the Kirindy-Mite National Park (KMNP), and the Mikea National Park (MIKEA). These three protected areas are characteristic of dry deciduous forest in Madagascar which is home to a unique floral and faunal biodiversity (Fig. 1). Many species are endemic to the region such as the symbolic *Adansonia grandidieri* of the Avenue of the Baobabs or *Microcebus berthae*, the smallest species of primate in the world. We derived historical forest cover change maps at 30 m resolution for the three study areas for the periods 1990–2000–2010–2017. We validated these maps with field observations. To identify the main causes of the deforestation, we conducted local household and environmental stakeholder surveys. A much clearer scheme emerged from this analysis explaining why environmental actions are currently failing to stop deforestation. Finally, based on a spatial deforestation model, we predicted the likely future of the tropical dry forest in 2050 in the three study areas in the absence of any efficient solution to curb deforestation.

**Figure 1:**
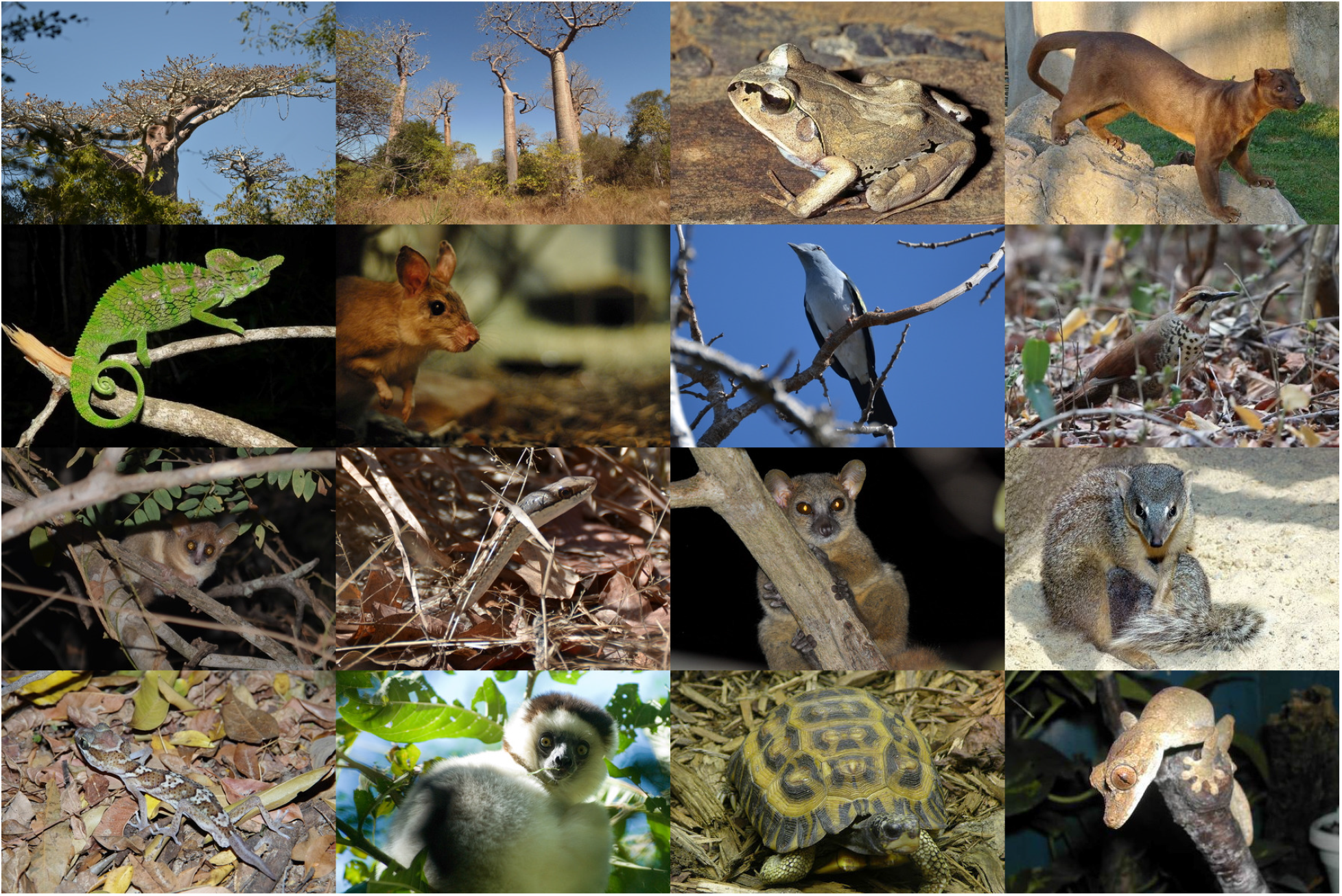
Emblematic species representative of the biodiversity in Western Madagascar. The dry forest of Western Madagascar is home to a very large number of species, many of which being endemic to the region. We present here some examples of this biodiversity for different taxonomic groups: Plants, Birds, Mammals (including Lemurs), Amphibians and Reptiles. From top-left to bottom-right: 1. *Adansonia grandidieri* (Baillon, 1888), 2. *Adansonia rubrostipa* (Jumelle & Perrier, 1909), 3. *Aglyptodactylus laticeps* (Glaw, Vences & Böhme, 1998) 4. *Cryptoprocta ferox* (Bennett, 1833), 5. *Furcifer labordi* (Grandidier, 1872), 6. *Hypogeomis antimena* (Grandidier, 1869), 7. *Leptosomus discolor* (Hermann, 1783), 8. *Mesitornis variegata* (Geoffroy Saint-Hilaire, 1838), 9. *Microcebus berthae* (Rasoloarison, Goodman & Ganzhorn, 2000), 10. *Mimophis mahafaliensis*, 11. *Mirza coquereli* (Grandidier, 1867), 12. *Mungotictis decemlineata* (Grandidier, 1867), 13. *Paroedura picta* (Peters, 1854), 14. *Propithecus verreauxi* (Grandidier, 1867), 15. *Pyxis planicauda* (Grandidier, 1867), 16. *Uroplatus guentheri* (Mocquard, 1908). Credits: 1,2,5,7,9,10,13,15: authors; 3: Miguel Vences; 11: Louise Jasper; 4,6,8,12,14,16: Wikipedia.

## 2 Materials and Methods

### 2.1 Historical deforestation maps

We used 30 m resolution cloud-free forest cover maps available for Madagascar for the years 1990, 2000, 2010, and 2017 (Vieilledent *et al*., 2018b,a). These maps were obtained combining the 1990 and 2000 forest cover maps by Harper *et al*. (2007) and the 2000–2017 tree cover loss data by Hansen *et al*. (2013). We used these national forest cover maps to derive forest cover change maps for the periods 1990–2000, 2000–2010, and 2010–2017 for our three study areas around the Menabe-Antimena New Protected Area (E: 44.20934, W: 44.81850, S: −20.36440, N: −19.55838, in decimal degrees), the Kirindy-Mite National Park (E: 43.67555, W: 44.35049, S: −21.38515, N: −20.50082), and the Mikea National Park (E: 43.11582, W: 44.17152, S: −22.98611, N: −21.62237).

For each period and each study area, we computed the mean annual deforested area (in ha/yr), and the mean annual deforestation rate *θ* (in %/yr) using the forest cover estimates *F*_1_ and *F*_2_ at dates *t*_1_ and *t*_2_ and the following formula (Puyravaud, 2003; Vieilledent *et al*., 2013): *θ* =1 – (1 – (*F*_1_ – *F*_2_)/*F*_1_)^(1/*T*)^, with *T* = *t*_2_ – *t*_1_ (in yr).

Field work was conducted during two weeks in March and June 2016 to verify the presence of the recent (2000–2010–2015) patches of deforestation in the three study areas. We validated the deforestation maps on these periods and identified the main causes of deforestation in the three study areas.

### 2.2 Deforestation model

Following Vieilledent *et al*. (2013), we considered two processes for modelling deforestation, a first one describing the intensity of deforestation (number of hectares of forest deforested each year) and a second one describing the location of the deforestation (spatial probability of deforestation). For the intensity of deforestation, we used two historical means at the study area level considering a conservative scenario (S1) and a worst-case scenario (S2). The conservative scenario assumed a lower deforestation rate corresponding to the mean annual deforestation observed in the period 2000–2010. The worst-case scenario assumed a higher deforestation rate corresponding to the mean annual deforestation observed in the period 2010–2017, during which the deforestation has dramatically increased.

For the location of the deforestation, we used a map of the spatial probability of deforestation at 30 m resolution for the year 2010 for Madagascar. The map is freely available at https://bioscenemada.cirad.fr/forestmaps. We used this national map for our three study areas (Appendix 1). The map was derived from a Binomial logistic regression model of deforestation where the observed deforestation at the pixel level in the period 2000–2010 was explained by several environmental spatial variables. Altitude, distance to forest edge, distance to main town, distance to main road, protected areas, and distance to past deforestation (period 1990–2000) were used as explanatory variables. These variables describe the accessibility, the land policy and the historical deforestation. The model also included spatial random effects at the regional scale (10 × 10 km grid cells) to account for the residual variability in the deforestation process which is not explained by the environmental variables (see https://ghislainv.github.io/forestatrisk for more details on model specifications).

To forecast the forest cover in 2050 for our three study areas, we removed the 2010 forest pixels with the highest probability of deforestation following the annual deforestation rate of the two intensity scenarios considered (conservative scenario S1 and worst-case scenario S2). As a rough validation of our projections, we computed the percentage of observed deforestation in the period 2010–2017 included in the predicted deforestation in the period 2010–2050.

### 2.3 Surveys

During two weeks in March and June 2016, we conducted surveys among local farmers and environmental stakeholders (Tab. 1) in the three study areas. The objectives of the surveys were firstly to identify the causes of the deforestation and secondly to assess the efficiency of the conservation actions implemented by the organisations in charge of the management of the protected areas, either the NGO Fanamby for MANAP or the parastatal association Madagascar National Parks for KMNP and MIKEA. Surveys were qualitative and included non-directive questions (no “yes-or-no” questions). Discussing with the “*Fokontany*” chief at Lambokely, we also obtained rough population estimates on the period 2010–2015 for the two villages of Kirindy and Lambokely. The “*Fokontany*” is a Malagasy administrative unit under the level of the township. Information from field surveys was cross-checked and complemented by a review of technical reports on the maize and peanut sectors in Madagascar (Youssi, 2008; Ministère de l’Agriculture, de l’Elevage et de la Pêche de Madagascar, 2004; Chan Mouie, 2016; Fauroux, 2000) and scientific studies on the deforestation process in south-east Madagascar (Scales, 2011; Zinner *et al*., 2014; Fauroux, 2000; Réau, 2002; Casse *et al*., 2004; Gardner *et al*., 2013). We also used the FAOSTAT website (http://www.fao.org/faostat) to obtain information on the national production of maize and peanut; and the UN Comtrade website (https://comtrade.un.org) to obtain information on the exportations.

**Table 1:** List of surveys conducted among local farmers and environmental stake-holders. Surveys were conducted around the Menabe-Antimena New Protected Area, the Kirindy-Mite National Park, and the Mikea National Park. The “*Fokontany*” is a Malagasy term defining an administrative territory under the level of the township. In Madagascar, the management of the protected areas is under the responsibility of Madagascar National Parks (MNP) or delegated to another environmental Non-Governmental Organisation (such as Fanamby for Menabe-Antimena New Protected Area).

| Id | Category | Name | Institution | Position | Date |
| --- | --- | --- | --- | --- | --- |
| 1 | National Parks | M. Toany | MNP | Director of the Mikea National Park | 15/03/16 |
| 2 | Migrant and farmer | Farmer 4 | Basibasy commune | Member of the Mikea community | 16/03/16 |
| 3 | Migrant and farmer | Farmers 5 and 6 | Antanimieva commune | Chiefs of family | 17/03/16 |
| 4 | Migrant and farmer | Farmers 7 and 8 | Ankantsakantsa commune | Chiefs of family | 18/03/16 |
| 5 | Administration | Mayor | Analamisampy commune | Mayor | 19/03/16 |
| 6 | Private Enterprise | Technicians | Ankililoaka commune | Agriculture and sales technicians | 20/03/16 |
| 7 | National Parks | Mamy Rakotobendrasana | MNP | Director of the conservation program for KMNP | 07/06/16 |
| 8 | Environmental NGO | Roland Eve | WWF | Landscape Planning and Management advisor for KMNP | 08/06/16 |
| 9 | Government representative | Mme Cynthia | DREF Menabe | Acting director of the Direction Régionale des Eaux et Forêt | 09/06/16 |
| 10 | Woman | Woman 1 | Kirindy village | Woman of farmer living in the Kirindy village | 10/06/16 |
| 11 | Migrants and farmers | Farmers 2 and 3 | Lambokely | Two family chiefs with women and children | 10/06/16 |
| 12 | Administration | Chief 1 | Lambokely | Fokontany chief | 10/06/16 |
| 13 | Environmental NGO | Tahina Riniavo | Fanamby | Director of the conservation program for MANAP | 10/06/16 |
| 14 | Migrant and farmer | Farmer 1 | Kirindy village | Chief of a family with a woman and eight children | 10/06/16 |

## 3 Results

### 3.1 Intensity and pattern of deforestation

Deforestation rates have continuously increased since 1990 for the MANAP and KMNP study areas (Tab. 2). After a decrease in 2000–2010, deforestation in the MIKEA study area has dramatically increased in 2010–2017 (Tab. 2). We estimated that 5429 ha, 4312 ha and 8210 ha of forest have disappeared annually in the period 2010–2017 in the MANAP, KMNP, and MIKEA study areas, respectively. This corresponds to annual deforestation rates of 2.16–4.31%/yr. Deforestation has more than doubled in the period 2010–2017 compared to the period 2000–2010, for which deforestation was estimated at 0.66–0.87%/yr.

**Table 2:**
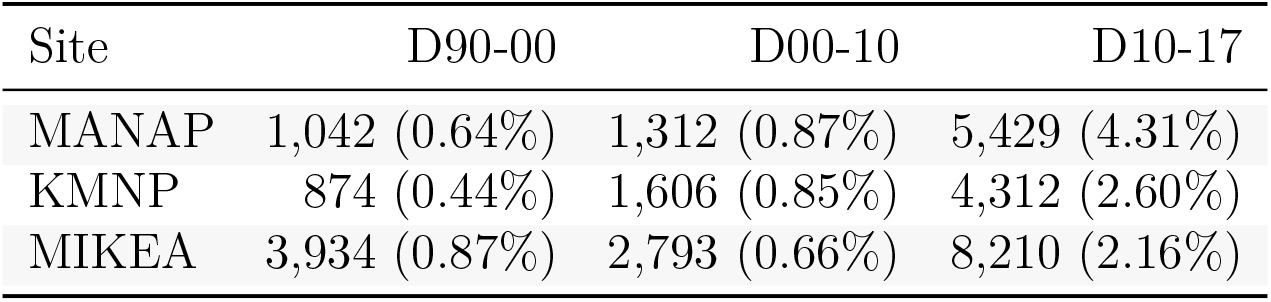
Change in annual deforestation with time. MANAP: Menabe-Antimena New Protected Area, KMNP: Kirindy-Mite National Park, MIKEA: Mikea National Park. *D90-00, D00-10*, and *D10-17*: annual deforestation (in ha/yr) for the periods 1990–2000, 2000–2010, and 2010–2017 respectively, followed by the annual deforestation rate in parenthesis (in %/yr). The annual deforestation has more than doubled on the period 2010–2017 compared to the period 2000–2010.

| Site | D90-00 | D00-10 | D10-17 |
| --- | --- | --- | --- |
| MANAP | 1,042 (0.64%) | 1,312 (0.87%) | 5,429 (4.31%) |
| KMNP | 874 (0.44%) | 1,606 (0.85%) | 4,312 (2.60%) |
| MIKEA | 3,934 (0.87%) | 2,793 (0.66%) | 8,210 (2.16%) |

In the MANAP study area, large patches of deforestation associated with slash-and-burn agriculture were identified around the Kirindy and Lambokely villages and to the south of Belo-sur-Tsiribihina town (Fig. 2a, label A).

**Figure 2:**
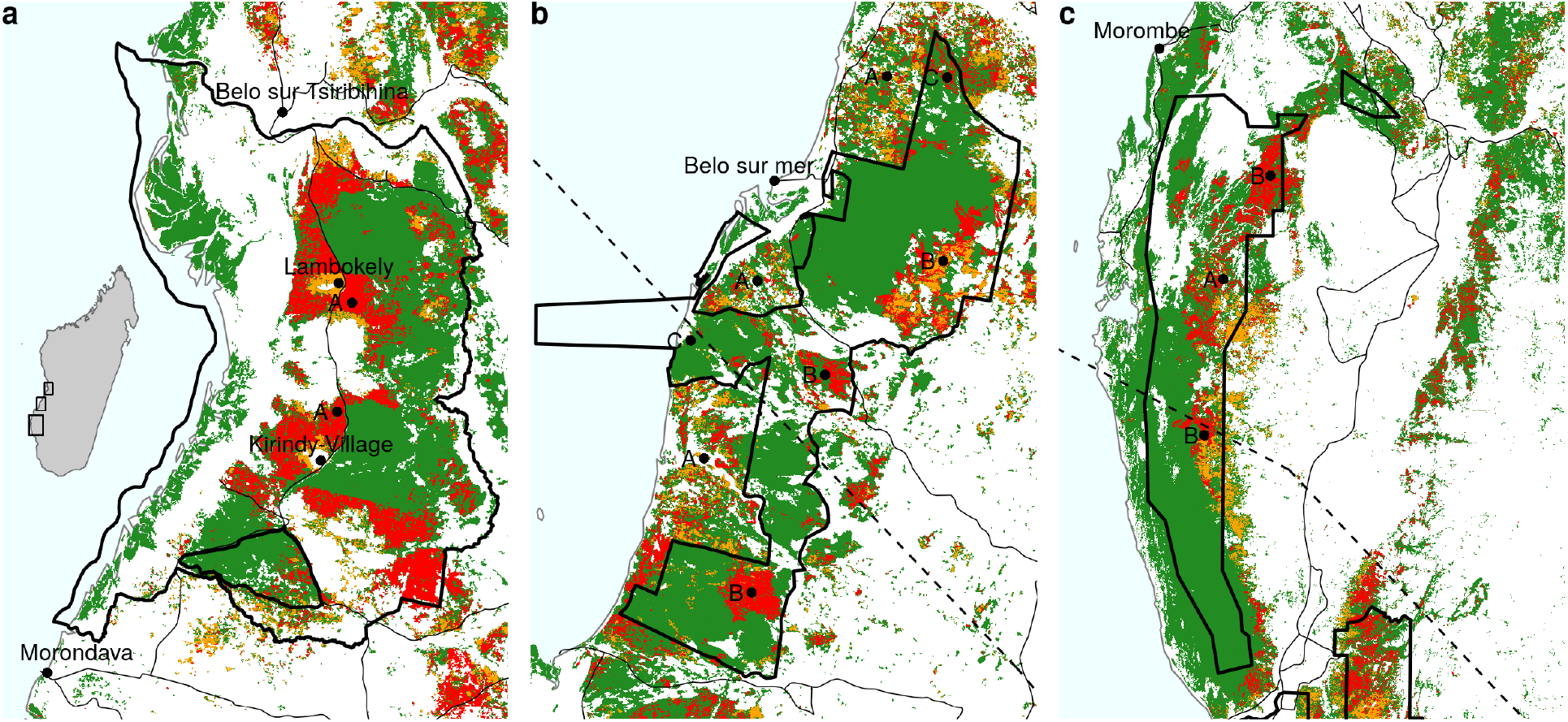
Historical deforestation on the period 2000–2010–2017 in the three study areas. Madagascar map is represented on the left panel (a), with MANAP study area at the north, KMNP study area in the middle, and MIKEA study area at the south (black rectangles). On each of the panels, the boundaries of the protected areas are represented with black polygones (source: Rebioma, http://rebioma.net). Main roads are represented with thin black lines (source: FTM BD500). Coast line is represented with a thin grey line. Morondava and Belo-sur-Tsiribihina are the main cities located near MANAP. Belo-sur-Mer is the main village located near KMNP. Morombe is the main village located near MIKEA. The dashed lines on b) and c) show the trajectories of the cyclones “*Fanele*” (January 2009) and “*Haruna*” (February 2013), respectively (source: JTWC, https://www.metoc.navy.mil/jtwc). Green: forest cover in 2017, orange: 2000–2010 deforestation, red: 2010-2017 deforestation (Vieilledent *et al*., 2018a). In the study areas, the main causes of deforestation are: (A) slash-and-burn agriculture (*“hatsake*”) for maize and peanut crops, (B) cyclones followed by uncontrolled fires, and (C) illegal logging.

In the KMNP study area, mosaic deforestation associated with slash-and-burn agriculture occurred outside the protected area, showing the relative effectiveness of the protected area to prevent deforestation in the short term (Fig. 2b, label A). Much larger patches of deforestation have been observed on the two periods 2005–2010 and 2010–2015 in the east part of the protected area (Fig. 2b, label B). These large patches of deforestation have been caused by the cyclone *Fanele* that occurred in January 2009 (Appendix 2) and which was followed by uncontrolled fires. Dispersed and small-scale deforestation has also been observed in the northern and western parts of the park associated with illegal logging activities (Fig. 2b, label C).

In the MIKEA study area, most of the deforestation was located in the north of the protected area (far from the Madagascar National Park office, which is located in the south of the study area) and was both due to slash and burn agriculture (Fig. 2c, label A) and to uncontrolled fires following cyclone *Haruna* which occurred in February 2013 (Fig. 2c, label B; and Appendix 2).

### 3.2 Deforestation drivers

#### 3.2.1 Proximate causes of deforestation: slash-and-burn agriculture and uncontrolled fires

In the MANAP study area, the main cause of deforestation was the slash-and-burn agriculture (locally known as “*hatsake*”) for maize (*Zea mays* L.) and peanut (*Arachis hypogaea* L.) crop (Fig. 3). The burning of forests allows expansion of cultivable areas and optimisation of labour productivity. It provides nutrient rich ash and light for crops, increasing yields and reducing the necessary time for weeding. In the KMNP and MIKEA study areas, slash-and-burn agriculture was also identified as a cause of deforestation, but preferentially outside protected areas (Fig. 2b-c, label A).

**Figure 3:**
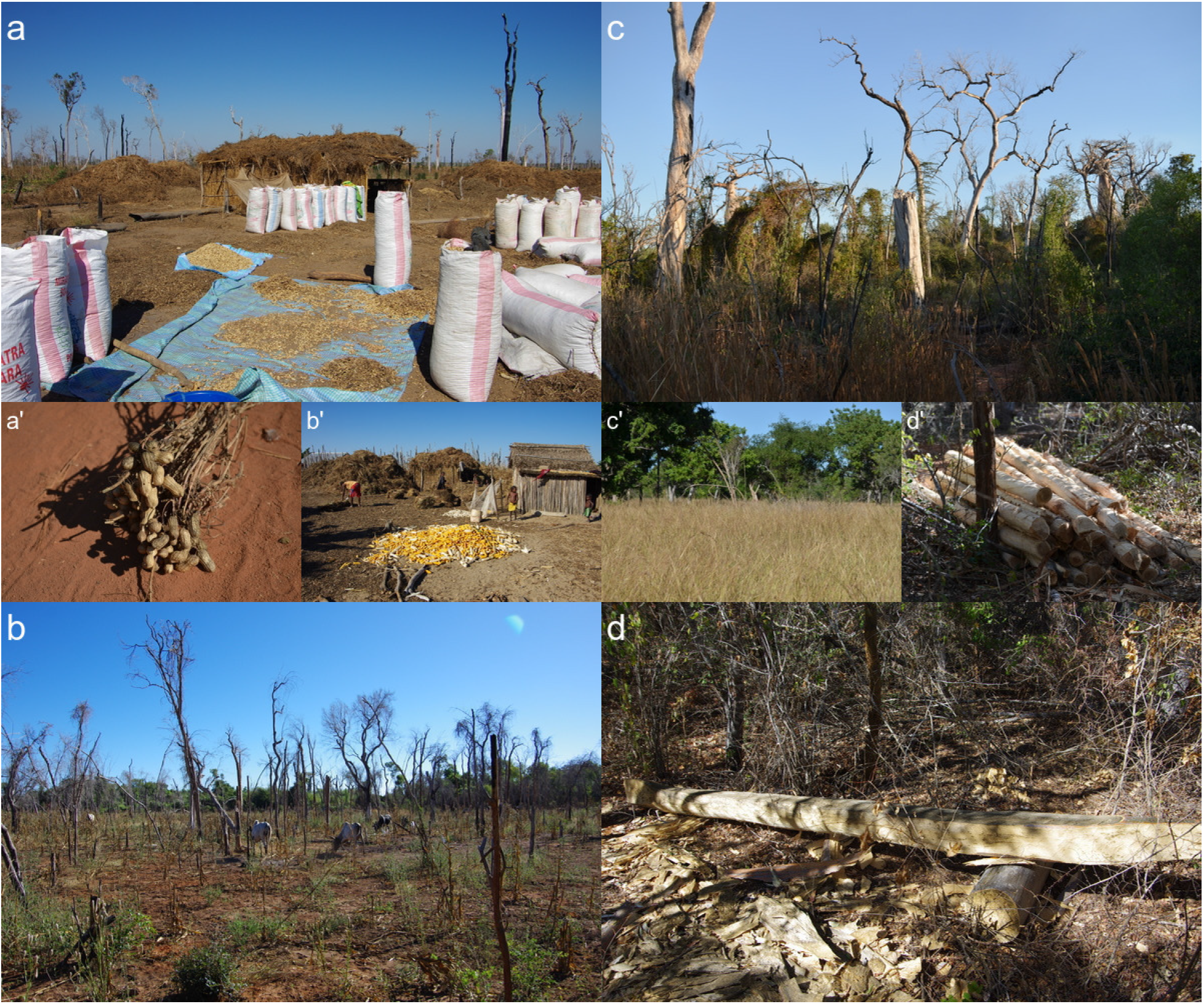
Main causes of deforestation in Western Madagascar. **a-a’**: *Slash-and-burn agriculture* (“hatsake”) *for peanut crop*. Peanut (a’) is cultivated as a cash crop. Most of the production is at the destination of the domestic market. **b-b’**: *Slash-and-burn agriculture for maize crop*. Maize (b’) is cultivated for auto-consumption and as a cash crop. The production of maize is at the destination of the domestic market and is used in particular for livestock feeding. **c-c’**: *Cyclone followed by uncontrolled fires*. Cyclone “*Fanele*” (2009) caused tree mortality and accumulation of wood fuel on the ground. As a consequence, uncontrolled fires set on nearby pastures (c’) spread over large areas of forest after 2009. **d-d’**: *Illegal logging*. Timbers are used for house and dugout canoe construction.

Inside the KMNP and MIKEA protected areas, uncontrolled fires (Fig. 3) were the main cause of deforestation (Fig. 2b-c, label B). People repeatedly set fire to former grasslands (called “*bozake*”) outside the protected area to obtain a flush of green pasture for their livestock. When uncontrolled, fires can spread across large areas of forest and cross the boundaries of the protected area. In 2009, the cyclone named *Fanele* (Appendix 2) impacted a large area of the forest in the Kirindy-Mitea National Park leaving a lot of wood fuel on the ground. This allowed uncontrolled fires to spread throughout large areas of the park in the years following the cyclone (Fig. 2b, label B). These fires were eventually stopped with water and sand by the agents of the park with the help of local villagers. The same thing happened in the MIKEA study area after cyclone *Haruna* in 2013 (Fig. 2c, label B and Appendix 2). Cyclones provide opportunities for local people to gain land on the forest. Following cyclones, it is more difficult to accuse people of destroying the forest and cyclones allow farmers to avoid the labour-intensive work of cutting down the trees before burning.

Illegal logging was also identified as a cause of forest degradation in the three study areas (Fig. 2b, label C). Illegal logging is not a direct cause of deforestation but a forest without precious wood is more easily burnt than an intact forest, and so degradation usually precedes deforestation. Timber (Fig. 3) is mainly used for house and boat construction and sold in local markets in Belo-sur-Mer and Morondava towns.

#### 3.2.2 Ultimate causes of deforestation

##### 3.2.2.1 Demographic growth and migration

The population of Kirindy and Lambokely villages (Fig. 2a) has been roughly multiplied by 5 between 2010 and 2015 (from about 600 to 3000 inhabitants for Kirindy and from about 1000 to 5000 for Lambokely). This increase was due both to demographic growth and migration. The demographic growth rate in Madagascar is close to 3%/yr (Vieilledent *et al*., 2013) which means that the population doubles every 25 years on average. In Lambokely and Kirindy villages, the families we surveyed had all more than six children. Also, the possibility of cropping cash crops have attracted many people from the south of Madagascar during the last years, in particular from the *Androy* and *Atsimo-Atsinanana* regions (South-East of Madagascar). First migrants arrived in the central Menabe to work in large agricultural concessions authorized by the French colonial government. Notably, many *Tandroy* migrants have arrived in the 1960s and established near the Beroboka village (located between Kirindy and Lambokely villages) to work in the sisal (*Agave sisalana* Perr.) plantation of the de Heaulme family which is now abandoned. Consecutive droughts and crop failure in South-Eastern Madagascar resulted in severe famines there, that forced several thousand *Tandroy* families to migrate to Western Madagascar in search for new farming land. Population increase in Western Madagascar has accentuated the pressures on forests.

##### 3.2.2.2 Cash crops and unregulated market

In the study areas we surveyed, peanuts (Fig. 3) was cultivated as a cash crop. Peanuts are consumed as whole seeds or transformed into peanut oil (Ministère de l’Agriculture, de l’Elevage et de la Pêche de Madagascar, 2004). According to FAOSTAT data (Tab. 3), the area harvested for peanut crop in Madagascar has increased of 50% in 7 years, from 52,000 ha in 2010 to 78,426 ha in 2017. At the same time, according to UN Comtrade data, peanut exports have also dramatically increased, from 1,233 T in 2010 to 27,370 T in 2017 (Tab. 3 and Appendix 3). In 2017, about half of the national peanut production was exported, mainly at the destination of the Asian market (Vietnam and Pakistan, in particular). During our stay in the field in June 2016, which took place in the middle of the peanut harvest, we observed an uninterrupted parade of trucks arriving empty in the villages of Lambokely and Kirindy (Fig. 2a) and leaving loaded with peanut bags.

**Table 3:** National statistics for maize and peanut production and exports. The area harvested is in hectares (ha), the yield is in tonne per ha per year (T/ha/yr), the total production and the exportation are in tonnes per year (T/yr). Sources: FAOSTAT (http://www.fao.org/faostat) for production and UN Comtrade (https://comtrade.un.org) for exports.

| Crop | Year | Area | Yield | Production | Exports |
| --- | --- | --- | --- | --- | --- |
| Maize | 2000 | 192,135 | 0.9 | 169,800 | 5,458 |
|  | 2005 | 252,838 | 1.5 | 391,000 | 13 |
|  | 2010 | 264,429 | 1.6 | 411,914 | 1,572 |
|  | 2015 | 194,269 | 1.7 | 329,367 | 4,458 |
|  | 2017 | 162,176 | 1.7 | 281,000 | 652 |
| Peanut | 2000 | 47,205 | 0.7 | 35,030 | 1,077 |
|  | 2005 | 54,506 | 0.7 | 38,053 | 659 |
|  | 2010 | 52,000 | 0.6 | 30,000 | 1,233 |
|  | 2015 | 66,591 | 0.9 | 58,483 | 17,716 |
|  | 2017 | 78,426 | 0.7 | 55,646 | 27,370 |

Our surveys indicated that maize (Fig. 3) was cultivated for auto-consumption (about 30% of the production) and as a cash crop (70%). FAOSTAT and UN Comtrade data indicate that only a small part of the national maize production was exported (about 5,000 T/yr out of a total production of about 200,000 T/yr), most part of the production being at the destination of the domestic market (Tab. 3 and Appendix 3). Several people we interviewed said that the maize production was sold to the Star company to brew the THB national beer. We computed that about 2,471 ha of maize are necessary to produce the annual volume of THB of 840,000 hL (Appendix 4). This is a relatively small area compared to the 249,186 ha of maize harvested annually in Madagascar (Tab. 3). Other sources (Ministère de l’Agriculture, de l’Elevage et de la Pêche de Madagascar, 2004) indicated that maize is used for livestock (poultry and pigs) feeding in Madagascar.

Farmers we interviewed sold the peanuts and maize at the price of 1,400 MGA (Madagascar Ariary) and 400 MGA per kilogram, respectively. For 2016, the production for a household was approximately of 1.6 T of peanut and 2.5 T of maize, thus providing an annual income of about 3.24 millions MGA. Farmers we interviewed said they invested the money earned from the sale of the maize and peanut harvest in zebu herd acquisition.

##### 3.2.2.3 Limits in the application of conservation policy

Since 1987 forest clearance has been illegal in Madagascar (Décret n°87-143, 20 April 1987), even outside the protected areas.

However, as underlined by NGO in charge of the protected area management, the law is not respected nor applied. Almost nobody is prosecuted for forest clearance. Small farmers can be prosecuted as examples but are rarely incarcerated. Large landholders or those inciting small farmers to clear the forest are not prosecuted. During our stay in the field, seven people were arrested for doing slash-and-burn agriculture but were released a few days later. The political crisis of 2009, followed by several years of political instability during which funding for protected area management was severely curtailed, has reinforced this state of lawlessness.

Several people we interviewed said that officials in the army or with political responsibilities, who are at the same time businessmen or entrepreneurs, are involved in this trade.

Moreover, authorities commonly have economic interests in not curbing deforestation as they are often involved in the trade associated with cash crops. Indeed, many politicians in Madagascar are also business leaders. Throughout the country, the Government has delegated management of the protected areas to external parties such as Fanamby and Madagascar National Parks. However, the structures in charge of the management of the protected areas have no legal enforcement powers. These powers are retained by the Malagasy Government which does not have the resources or the will to implement them adequately. Thus, the role of protected area managers is limited to raising awareness about forest conservation issues, inventorying and monitoring the biodiversity in the parks, and organizing community and NGO patrols to discourage forest clearance or report offences. They do not have the right to arrest people or to draw up a report and decide on a fine. NGOs also engage local people as conservation partners (in some cases named “*polis ny ala*”) to try to make them stewards of their forest but these have practically no power and social pressures mean that they are often unwilling to report illegal acts committed by neighbours or relatives.

### 3.3 Projected deforestation

Following the conservative scenario S1 (projecting 2000–2010 mean annual deforestation) and the worst-case scenario S2 (projecting 2010–2017 mean annual deforestation), we predicted that 38-93% of the forest present in 2000 will have disappeared in 2050 (Tab. 4). In the period 2000–2017, around 25% of the forest has already disappeared. Forest in 2050 should remain predominantly in the protected areas but deforestation is not expected to stop at the boundaries of the parks (Fig. 4). The model predicted that deforestation in the future should occur close to places were deforestation occurred in the past, thus correctly simulating the contagious process of deforestation (Fig. 4). Deforestation is also more likely to occur at short distances to villages and roads and at the forest edge (Fig. 4). Forest fragmentation is also predicted to increase in association with deforestation (higher number of disconnected forest patches in Fig. 4). Most of the deforestation observed in the period 2010–2017 was included in the deforested area predicted by the model in the period 2010–2050 (47-100% for MANAP, 56-100% for KMNP, and 43-88% for MIKEA for scenarios S1 and S2, respectively), thus validating partly the predictions regarding the location of the future deforestation.

**Table 4:** Change in forest cover with time. MANAP: Menabe-Antimena New Protected Area, KMNP: Kirindy-Mite National Park, MIKEA: Mikea National Park. *Area:* land area (in ha). *F2000, F2010* and *F2017*: forest area (in ha) for the years 2000, 2010 and 2017, respectively. *F2050*: projected forest area (in ha) for the year 2050. About 25% of the forest have disappeared on the period 2000–2017 in the three sites and we predict the loss of 38-93% of the forest on the period 2000–2050 for the three sites assuming a conservative scenario S1 (projecting the 2000–2010 annual deforestation) or a worst-case scenario S2 (projecting the 2010–2017 annual deforestation), respectively.

| Site | Area | F1990 | F2000 | F2010 | F2017 | F2050,S1 | F2050,S2 |
| --- | --- | --- | --- | --- | --- | --- | --- |
| MANAP | 349,360 | 166,648 | 156,224 | 143,108 | 105,109 | 90,644 | 0 |
| KMNP | 403,080 | 203,920 | 195,175 | 179,117 | 148,932 | 114,887 | 6,637 |
| MIKEA | 1,208,910 | 472,542 | 433,199 | 405,274 | 347,804 | 293,575 | 76,878 |

**Figure 4:**
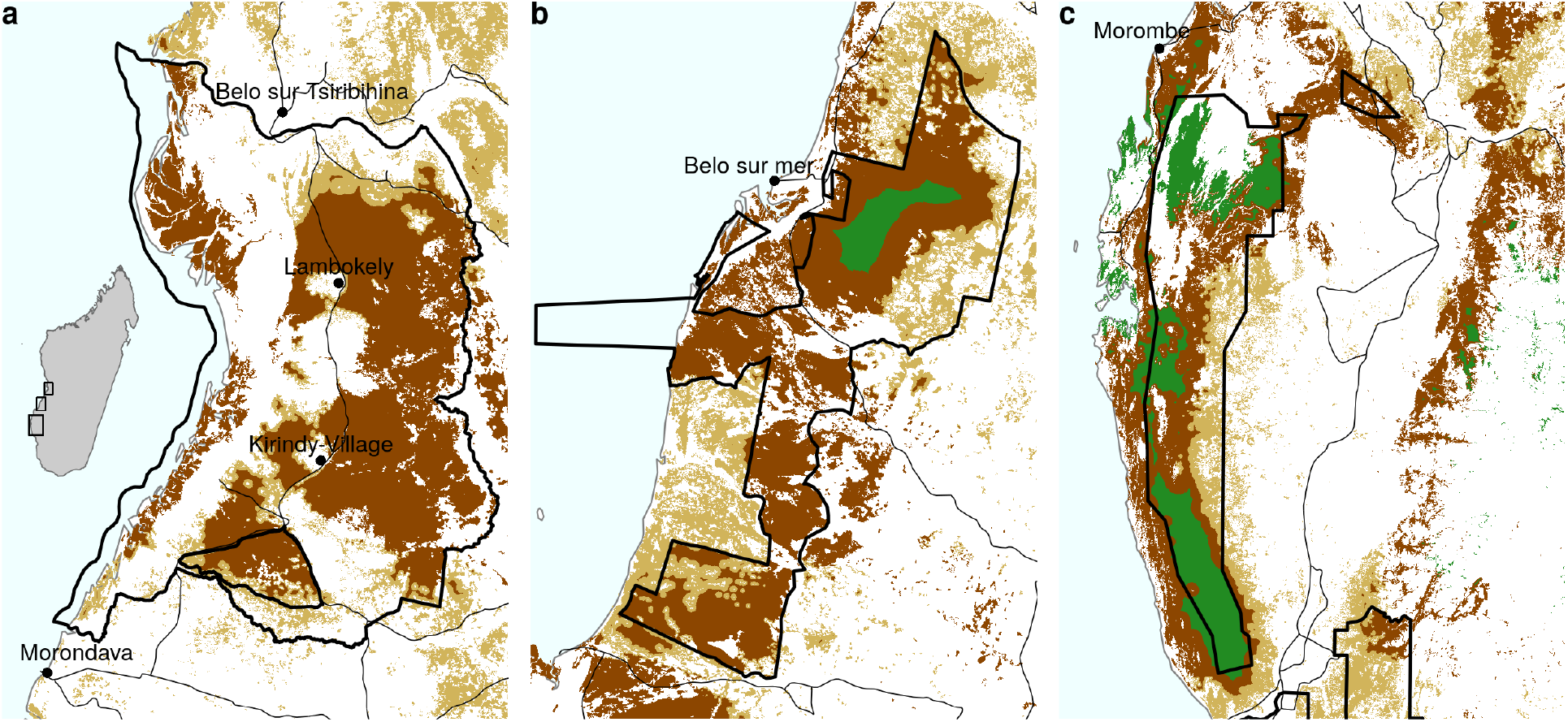
Projected deforestation on the period 2010–2050. Green: projected forest cover in 2050, light brown: 2010–2050 deforestation following conservative scenario S1 (projecting 2000–2010 mean annual deforestation), dark brown: 2010–2050 additional deforestation following scenario S2 (projecting 2010–2017 mean annual deforestation). Most of the 2010–2017 deforestation observed on Fig. 2 is included in the 2010–2050 projections. We predicted a loss of 38-93% of the forest cover in 2050 compared to 2000 following scenario S1 and S2, respectively. Remaining forest in 2050 under scenario S1 should be concentrated inside protected areas. We predicted a complete loss of the forest for the MANAP study area following scenario S2.

## 4 Discussion

### 4.1 Historical deforestation, projections and consequences on livelihoods, biodiversity and climate-change

For the three study areas, we have shown a strong increase of the annual deforestation rates on the period 1990–2015. This increase, also observed by Scales (2011) and Zinner *et al*. (2014) in the region, clearly demonstrate the inefficiency of the recent environmental policies to reduce deforestation. In the absence of any efficient future policy to curb deforestation, we predicted a 36–59% forest loss on the period 2000–2050. This scenario, confirmed by the results of Zinner *et al*. (2014), would be terrible for both local villagers in terms of livelihoods and at the global scale for biodiversity and carbon emissions. In term of loss of biodiversity, Allnutt *et al*. (2008) estimated a 9% decrease in the number of species after a deforestation of 40% on the period 1950–2000. In our case, given a forest loss of 36–67%, we can assume that the biodiversity loss would be of the same order of magnitude on the period 2000–2050. Many species endemic to the region, such as *Hypogeomis antimena, Mungotictis decemlineata, Microcebus berthae* and *Pyxis planicauda* could experience a dramatic demise of their populations resulting in a rapid tailspin toward extinction. Dry forests sequester an average of 52 Mg/ha (1Mg=10^6^g) (Vieilledent *et al*., 2016). Considering the predicted forest cover loss of 229,384–393,119 ha on the period 2010–2050 for the three study areas combined, deforestation would lead to emissions of 11.9–20.4 Tg (1Tg=10^12^g) of carbon in the atmosphere, thus contributing significantly to climate change.

### 4.2 Deforestation is not associated directly to poverty but to unregulated market and bad governance

Our field observations and surveys indicated that a large part of the deforestation was attributable to slash-and-burn agriculture for cash crops (maize and peanut). The major part of the maize and peanut production is at the destination of the domestic market.

Increase in area harvested for peanut crop is clearly associated to an increase in peanut exports toward the Asian market.

While a large part of the maize production in Western Madagascar have been exported through the ports of Morondava and Tulear during the successive maize booms in the 20th century (Scales, 2011; Fauroux, 2000), the production of maize seems to be presently mainly at the destination of the domestic market.

Area harvested for maize crop are relatively constant since 2000, and slightly decreasing since 2010.

In particular, maize is used for livestock feeding. The annual household income we computed associated to the selling of cash crops (about 3.24 millions MGA) was relatively high compared to the estimated median household income of 2.02 millions MGA (Gallup World Poll from the Gallup Organization). With the money earned from the sale of the maize and peanut harvest, farmers invest in accumulating zebu herd (Casse *et al*., 2004; Réau, 2002). Buying zebu herd is a way for farmers of saving money such as a bank would do it (Réau, 2002). In Madagascar, cattle also represent status, wealth, and cultural identity (Hobbs, 2016). In Western Madagascar, people’s diet is mainly composed of manioc (*Manihot esculenta* (Crantz)), wild and cultivated yams (*Dioscorea spp*.), and maize (Falola & Jean-Jacques, 2015). The main purpose of the slash-and-burn agriculture is thus not to obtain food for subsistence but to cultivate cash crop in order to invest in livestock acquisition and to secure new farming lands (Fauroux, 2000). In line with other studies (Grandin *et al*., 1988; Jarosz, 1993; Scales, 2011; Gardner *et al*., 2013), we underline that it would be over simplistic to reduce the causes of the deforestation to the poverty of local population.

Of course, Malagasy farmers do not make large profits and do not harvest most of the benefits associated to the culture of maize and peanut. Many intermediaries, such as storers, domestic transporters, resellers, exporters and corrupt officials reap most of the profits. Foreign entrepreneurs (European and Chinese) as well as entrepreneurs from the Malagasy elite (including “*Karana*”, the descendants of Indo-Pakistani migrants) connect rural households to the domestic market. Field surveys reported that several officials in the army or with political responsibilities, who are at the same time businessmen or entrepreneurs, are involved in this trade. As a consequence, stopping deforestation would go against the economic interest of some people having decisional and political power in Madagascar. In previous studies, Jarosz (1993) and Scales (2011) have clearly shown how economic booms of agricultural commodities and policies have driven deforestation in Madagascar during French colonization. Nowadays, deforestation in central Menabe is driven by the increasing demand in maize and peanut on the domestic market and by the possibility of making economic profits in these sectors.

Slash-and-burn agriculture for cash crops and the economic profits are made possible by the non-application of the environmental law in Madagascar. Forest clearance is illegal in Madagascar since 1987 (Décret n°87-143, 20 April 1987). In the field, the agents of MNP and Fanamby have no legal enforcement powers to arrest offender even when apprehended. Under-resourced local Government agencies that retain such legal powers are generally corrupt and have a low technical capacity, making their actions virtually ineffective. As a consequence, illegal activities such as slash-and-burn agriculture or setting fire to open pastures continue in full sight of everyone. This situation has been accentuated by the political crisis of 2009-2012 in Madagascar (Ploch & Cook, 2012). Madagascar is a fragile State country (the Fund for Peace, 2016). Common characteristics of a fragile State include a central Government so weak or ineffective that it has little practical control over much of its territory. Our study confirms that the deforestation problem in Madagascar is more a governance problem in a context of unregulated economy than an economic development problem. Interestingly, two global studies have recently shown that corruption (Venter *et al*., 2016), overexploitation and agriculture (including crop farming and livestock farming) (Maxwell *et al*., 2016) were responsible for a major part of the biodiversity loss, thus reflecting our results.

### 4.3 Enforcing law and controlling agricultural sector to stop deforestation in Western Madagascar

If we want to stop deforestation in western Madagascar, it seems necessary to rethink the conservation strategies we adopted during the last 30 years, which was based on extended the protected area network and alleviating poverty. One solution we see would be that national companies engage into ecological certification for agriculture commodities in Madagascar (Laurance *et al*., 2010; Kiker & Putz, 1997). Relative to estimates of conservation costs in the developing world, it has been shown that existing levels of environmental aid are insufficient (Miller *et al*., 2013). If the amount of the environmental aid have to be increased, the targets have also to be redefined and improving governance in Madagascar should be considered a priority (Smith *et al*., 2003). Brazil, which have reduced by two third the Amazon deforestation on 2005–2011 compared to 1996–2005, is a good example to show that both law enforcement (enforcement of the Brazilian Forest code) and control of the agricultural sector (voluntary moratoriums by the soy bean and beef industries) are efficient ways to rapidly curb deforestation (Boucher *et al*., 2013; Nepstad *et al*., 2009). This success, even if temporary, should inspire conservationists working in Madagascar and in other developing countries facing tropical deforestation in general.

Following the recent election of Andry Rajoelina as president of Madagascar, several researchers and conservationists have recently propose solutions to takle the decline in the rule of law in Madagascar (Jones *et al*., 2019). Among the solutions they propose, they suggest in particular to Tackle environmental crime and Invest in Madagascar’s Protected Areas. Out study demonstrates that…

## 5 Acknowledgements

This study took form in the framework of the BioSceneMada project (https://bioscenemada.cirad.fr) and the Roadless forests project (https://forobs.jrc.ec.europa.eu/roadless). The BioSceneMada project was funded by FRB (Fondation pour la Recherche sur la Biodiversité) and the FFEM (Fond Français pour l’Environnement Mondial) under the project agreement AAP-SCEN-2013 I. The Roadless forests project was funded by the European Commission. We thank all the people who welcomed us and kindly provided us with useful information during our field trip, both institutional people (from DREF, MNP, WWF and Fanamby) and villagers. The authors declare no conflict of interest.

## 6 Data availability statement

All the data and codes used for this study are made available publicly in the `menabe` repository on the GitHub platform at the following web address: https://github.com/ghislainv/menabe.git. The results and the manuscript are fully reproducible running the R script `menabe.R` from the `menabe` git repository.

## 9 Appendices

### 9.1 Appendix 1: Spatial probability of deforestation

**Figure A1:**
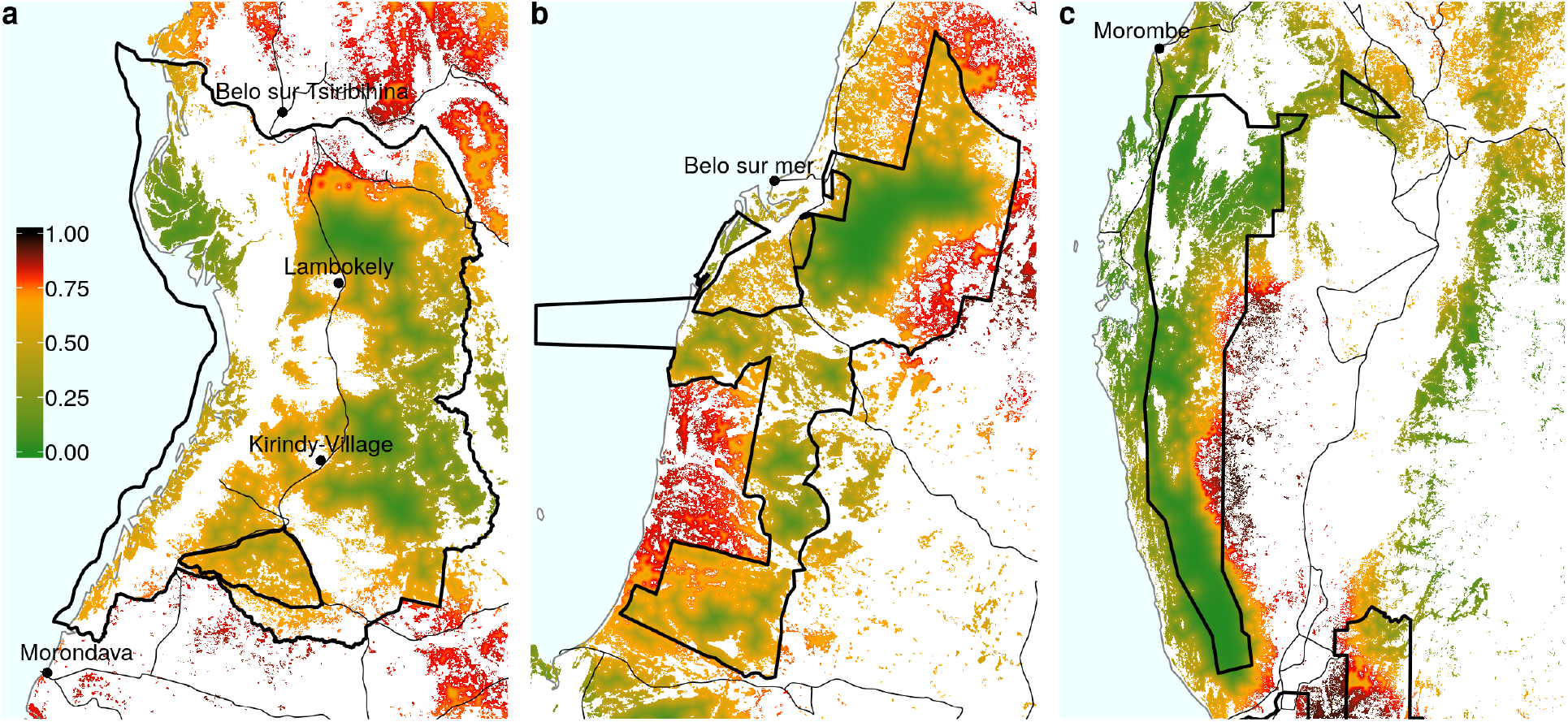
Spatial probability of deforestation for the year 2010. Source: https://bioscenemada.cirad.fr/forestmaps. The spatial probability of deforestation is modelled as a function of the altitude, the distance to forest edge, the distance to main town, the distance to main road, the protected areas, and the distance to past deforestation. These variables describe the accessibility, the land policy and the historical deforestation. The deforestation model also includes spatial random effects at the regional scale to account for the residual variability in the deforestation process which is not explained by the environmental variables (see https://ghislainv.github.io/forestatrisk for more details on model specifications).

### 9.2 Appendix 2: *Fanele* and *Haruna* cyclones

**Figure A2:**
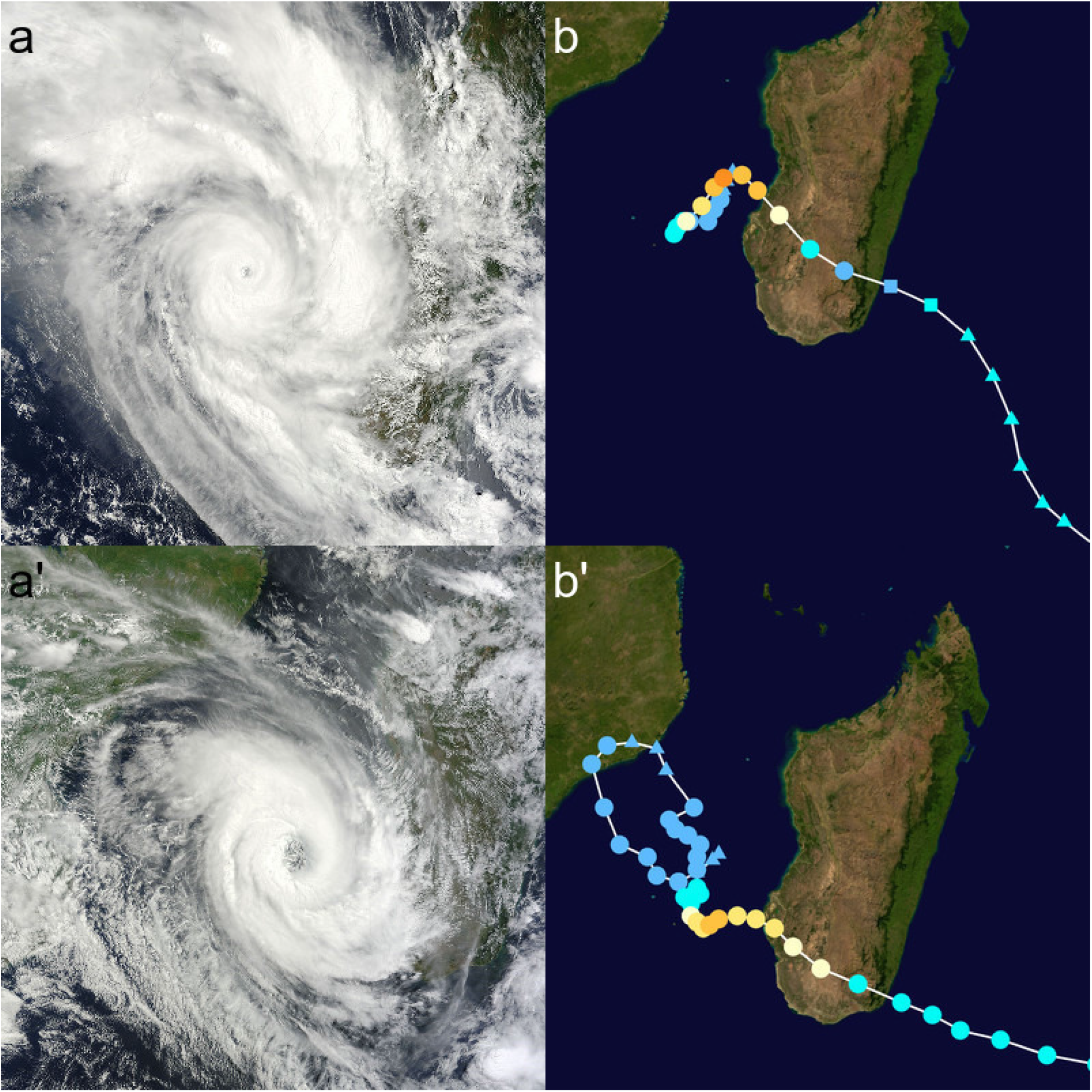
Satellite images and trajectories of cyclones *Fanele* and *Haruna* over Madagascar. **a-b**: Cyclone *Fanele* in January 2009. **a’-b’**: Cyclone *Haruna* in February 2013. **a-a’**: MODIS satellite image of the cyclone near peak intensity. **b-b’**: Track and intensity of the cyclone, according to the Saffir-Simpson scale. Highest winds for *Fanele:* 10–min sustained = 185 km/h, 1–min sustained = 215 km/h, gusts = 260 km/h. Highest winds for *Haruna:* 10–min sustained = 150 km/h, 1–min sustained = 195 km/h, gusts = not available. Source: Wikipedia, https://en.wikipedia.org/wiki/Cyclone_Fanele, https://en.wikipedia.org/wiki/Cyclone_Haruna.

### 9.3 Appendix 3: Change in maize and peanut exports

**Figure A3:**
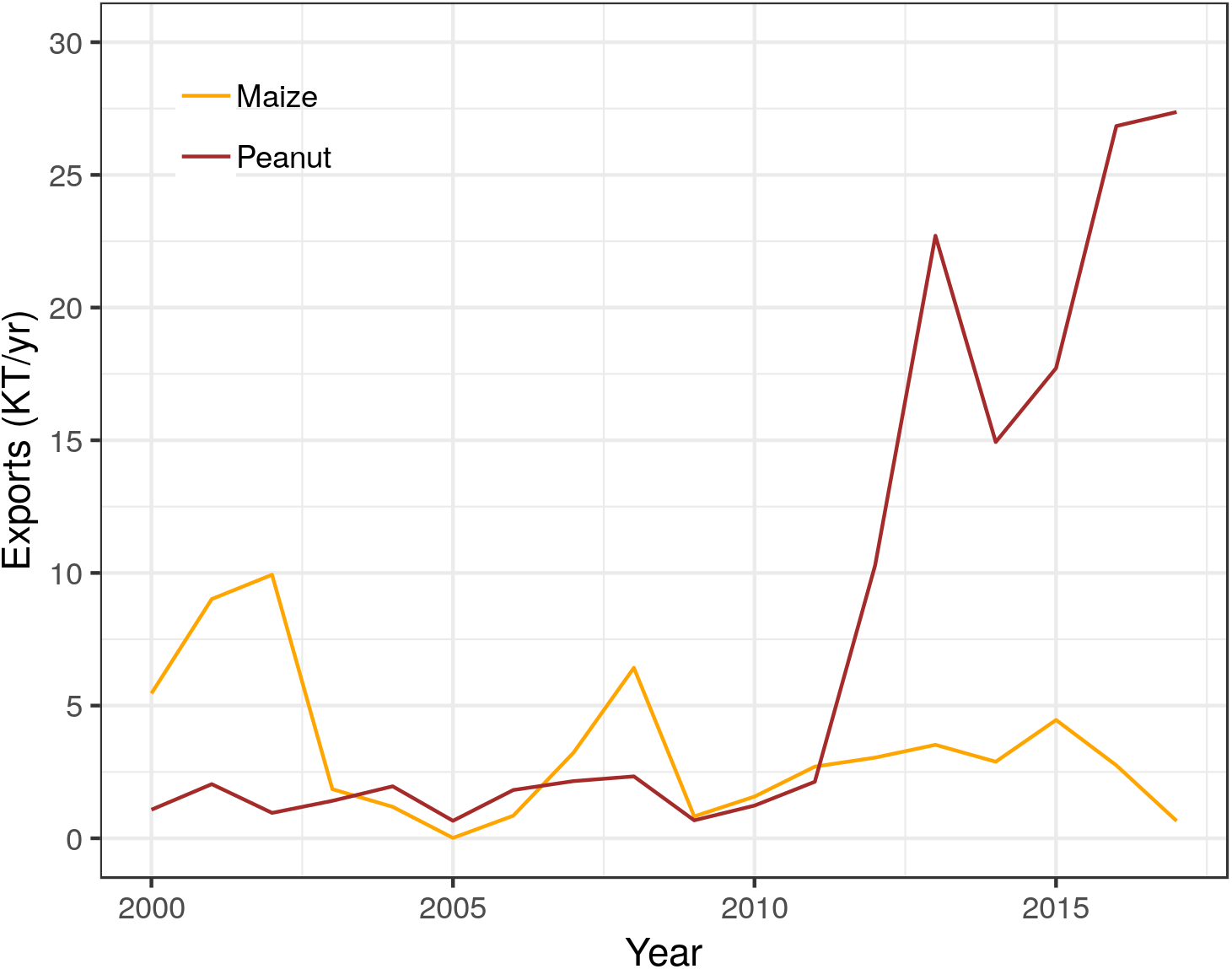
Annual exports in the period 2000–2017 for maize and peanut crops. Exports are reported at the national level for Madagascar in KT/yr. While maize exports are stable around 5 KT/yr in the period 2000–2017, peanut exports have boomed since 2011, with about 2.5 KT exported in 2011 to about 27,500 KT in 2017. Sources: UN Comtrade (https://comtrade.un.org).

### 9.4 Appendix 4: Maize and beer consumption in Madagascar

Three Horses Beer (better known locally as THB) has been brewed by Star Breweries of Madagascar since 1958. It is the highest selling beer in Madagascar and has been described as emblematic of the country. THB is sold nationwide and since 2005 has been exported to such markets as France, Reunion Island, Comoros and Mayotte. THB Pilsener, the most common variant of THB, is produced from mostly local barley, corn and hops. The mash is a blend of malt (sprouted barley) and corn in an 80/20 ratio to which water and hops are added. In 2006, the Star Brewery was producing 700,000 hectoliters (hL) of beer annually. Star Breweries invested over four million euros into improving its factories in the 2009–2011 period, resulting in a 20% increase in production capacity (source: Wikipedia, https://en.wikipedia.org/wiki/Three_Horses_Beer). As a consequence, we can assume that about 840,000 hL of THB Pilsener have been produced each year since 2011.

We assumed that about 1.25 kg of grain is needed to produce 1 kg of malt, and about 20 kg of malt are needed to produce 1 hL of beer (Kreisz, 2009). For a total of 840,000 hL of beer, 1.25 × 20 × 840,000 / 1000 = 21,000 tonnes (T) of grain are needed. Because the mash for the THB Pilsener is a blend of malt and corn in an 80/20 ratio, about 4,200 T of maize grain are necessary to produce the annual 840,000 hL of beer.

Given the 2010–2017 average yield of 1.7 T/ha for maize crop in Madagascar (Tab. 3), about 2,471 ha of maize crop are necessary to produce the local beer. This number is very small compared to the 225,084 ha of maize harvested at the national level (Tab. 3).

